# Crystal structure of a thermophilic family II inorganic pyrophosphatase enabling high-temperature adaptation in *Thermodesulfobacterium commune*

**DOI:** 10.1101/2025.02.25.640086

**Authors:** Saki Maruoka, Takamasa Teramoto, Keiichi Watanabe, Yoshimitsu Kakuta

## Abstract

Inorganic pyrophosphatases (PPases) are crucial for energy metabolism and classified into families with distinct metal ion requirements and structural features. This study is the first to successfully express, purify, and structurally characterize a thermophilic family II PPase isolated from *Thermodesulfobacterium commune* (*Tc*PPase). *Tc*PPase, which is optimally activated by Co^2+^ and Mn^2+^, required both the N- and C-terminal domains for complete catalytic function. Comparative structural analyses of psychrophilic and mesophilic homologs revealed that the enhanced thermal stability of *Tc*PPase was due to its strong hydrophobic interactions, high proline content, dense hydrogen bond network, and additional salt bridges. These findings reveal the molecular basis for the thermal adaptation of family II PPases, providing valuable insights for thermostable enzyme engineering for biotechnological applications.

## Introduction

Inorganic pyrophosphatases (PPases) catalyze the hydrolysis of inorganic pyrophosphate (PPi) into two phosphate (Pi) molecules, and this reaction is critical for several biosynthetic pathways and cellular energy homeostasis [1,2]. Specifically, family II PPases require divalent metal ions (e.g., Mg^2+^, Mn^2+^, and Co^2+^) and exhibit distinct structural and mechanistic features [3,4].

Thermal adaptation of enzymes often involves multiple structural strategies, such as enhanced hydrophobic packing, increased hydrogen bond networks, and additional salt bridges, which collectively stabilize the protein under extreme conditions [5–13]. Psychrophilic and mesophilic family II PPases have been extensively studied [14,15]; however, thermophilic family II PPases have not been properly characterized, with the molecular basis of their remarkable stability and catalytic efficiency at high temperatures remaining unclear.

In this study, we focused on a thermophilic family II PPase of *Thermodesulfobacterium commune* (*Tc*PPase), which thrives at a growth temperature of 70 °C [16]. We successfully expressed purified *Tc*PPase and determined its crystal structure. Biochemical characterization revealed that *Tc*PPase achieved maximal activity in the presence of Co^2+^ and Mn^2+^ and required both its N-terminal DHH (Asp-His-His) and C-terminal DHHA2 (Asp-His-His-associated type 2) domains for catalytic function. To elucidate its thermal adaptation mechanisms, we compared *Tc*PPase with psychrophilic family II PPases to determine the structural features responsible for enzymatic stability and proper function at elevated temperatures.

This study provides valuable insights into the structural and biochemical strategies used by family II PPases to enhance their thermal stability. Additionally, it provides a basis for the rational design of new thermostable biocatalysts with enhanced industrial and biotechnological applications.

## Materials and Methods

### Expression and purification of metal-free TcPPase

A cDNA encoding full-length *Tc*PPase (residues 1–320) was subcloned into the pE_SUMO vector (Life Sensors, Malvern, PA, USA) to introduce an N-terminal His_6_-SUMO tag. The recombinant protein was expressed in *Escherichia coli* BL21-CodonPlus (DE3)-RIL (Agilent Technologies, Santa Clara, CA, USA) via induction with 0.5 mM isopropyl-β-D-thiogalactoside at 20 °C for 16 h. After harvesting via centrifugation (2,500g, 30 min, 20°C), the cells were resuspended in a lysis buffer (50 mM Tris-HCl [pH 8.0] and 500 mM NaCl) and stored at −80 °C until use. The cells were lysed via sonication, and the lysates were clarified via centrifugation. The soluble fraction was further applied to an Ni-NTA agarose column (Fujifilm Wako, Osaka, Japan) equilibrated with the lysis buffer containing 20 mM imidazole. His_6_-SUMO fusion protein was eluted using the same buffer supplemented with 400 mM imidazole. After overnight cleavage with 0.2 mg of Ulp1 protease, the mixture was dialyzed against 50 mM Tris-HCl (pH 7.5), 500 mM NaCl, and 20 mM EDTA. Final purification was achieved using the Superdex 200 16/60 pg column (Cytiva, Chicago, IL, USA) equilibrated with 50 mM Tris-HCl (pH 7.5), 500 mM NaCl, and 20 µM EDTA. The purified protein was concentrated to 20 mg/mL.

### Preparation of metal-activated TcPPase

To prepare metal-activated *Tc*PPase, 0.5 mg/mL of metal-free *Tc*PPase was incubated with 2.5 mM divalent cations (Mg^2+^, Mn^2+^, Co^2+^, Ni^2+^, Ca^2+^, and Zn^2+^) at 20 °C for 2 h in a buffer containing 100 mM Tris-HCl (pH 7.5), 50 mM NaCl, and 20 µM EDTA.

### Expression and purification of the TcPPase N-terminal domain

A cDNA fragment encoding the N-terminal domain of *Tc*PPase (residues 1–200) was subcloned into the pE_SUMO vector (Life Sensors) to introduce the N-terminal His_6_-SUMO tag. The construct was generated using the Takara PrimeSTAR Mutagenesis Basal Kit (Takara Bio, Shiga, Japan), with pE_SUMO TcPPase as the template. N-terminal DHH domain-specific primers were synthesized by Hokkaido System Science (Sapporo, Japan). Expression and purification of the N-terminal domain were performed using the same procedure described for the full-length metal-free *Tc*PPase.

### Enzyme activity and kinetic analyses

Enzyme activity was measured using the molybdenum blue method. A 10-µL aliquot of *Tc*PPase was mixed with 110 µL of 1 mM potassium pyrophosphate (K_4_PPi) in 100 mM Tris-HCl (pH 7.5), 50 mM NaCl, and 5 mM MgCl_2_ and incubated at 25 °C for 3 min. The reaction was quenched by adding 30 µL of 50 mM H_2_SO_4_. To develop color, 150 µL of 1% ammonium molybdate in 0.05% K_2_SO_4_ and 1% sodium ascorbate was added. After 40 min, the liberated phosphate was quantified at 750 nm using a microplate reader (Nivo S; PerkinElmer, Waltham, MA, USA) and standard phosphate calibration curve (0–400 µM phosphate). Specific activity (U/mg) was defined as 1 μmol of PPi hydrolyzed per min per mg of protein. One unit of activity corresponded to the formation of 2 µmol of phosphate per min from 1 µmol of PPi under the assay conditions.

*K*_m_ and *k*_cat_ values were determined from the velocity data at various substrate concentrations using the GraphPad Prism 5 (GraphPad Software Inc., San Diego, CA.). PPi concentration ranged from 0 to 800 µM.

### Crystallization, data collection, structure determination, and refinement

Crystals of Mn^2+^-*Tc*PPase were obtained via sitting-drop vapor diffusion at 20 °C. Drops were prepared by mixing 200 nL of protein solution (34 mg/mL) containing 50 mM Tris-HCl (pH 7.5), 500 mM NaCl, 20 µM EDTA, 5 mM NaF, 5 mM imidodiphosphate sodium salt (PNP), and 10 mM MgCl_2_ with 200 nL of reservoir solution containing 0.2 M sodium citrate tribasic dihydrate and 20% (w/v) PEG 3350 at pH 8.3. The crystals were cryoprotected using the reservoir solution containing 20% glycerol and flash-frozen in liquid nitrogen.

X-ray diffraction data were collected at BL45XU (SPring-8, Hyogo, Japan) at a wavelength of 1.000 Å. Data was processed using the ZOO system [17]. Initial phases were determined via molecular replacement with Phaser [18] using *Sh*PPase (Protein Data Bank [PDB]: 6LL7) [14] as a search model. Model building and refinement were performed using Coot [19] and phenix.refine [20]. Final structures were validated using MolProbity [21], and the results are summarized in Table S1. The atomic coordinate and structure factor of the crystal structure have been deposited in the Protein Data Bank (PDB) under accession code 9M0M.

### Structural and surface analyses

Surface properties and secondary structures of *Tc*PPase, *Bacillus subtilis* family II PPase (*Bs*PPase), and *Shewanella* sp. AS-11 family II PPase (*Sh*PPase) were analyzed using MOE (version 2023.09; Chemical Computing Group, Montreal, QC, Canada). Hydrophobic/hydrophilic surface areas were computed from the atomic properties and solvent accessibility. Accessible surface area was calculated using a 1.4-Å probe radius. The residues in the accessible surface area at 0 Å were considered as buried residues. Then, secondary structure content, including the helix and strand percentages, was determined using the DSSP algorithm in MOE. Protein structures were energy-minimized using the Amber99 force field with default parameters. All calculations followed the standard protocols of MOE, unless stated otherwise.

## Results

### Divalent cation requirements and temperature effects

Recombinant full-length *Tc*PPase and isolated N-terminal DHH domain were successfully purified. Under various temperatures (20–80 °C), full-length *Tc*PPase showed optimal activity at 40 °C, irrespective of the divalent cation (Fig. 1A). Among the tested cations, highest activity was observed with Co^2+^ and Mn^2+^, followed by Ca^2+^, Ni^2+^, Mg^2+^, and Zn^2+^ (Fig. 1A), consistent with previously reported preferences for family II PPases [22,23]. Unlike psychrophilic *Sh*PPase, temperature-dependent activity of *Tc*PPase does not differ significantly with different metal ions [23,24]. Here, the isolated N-terminal DHH domain showed no detectable activity, confirming the need for both domains for its catalytic activity.

**Fig. 1.**
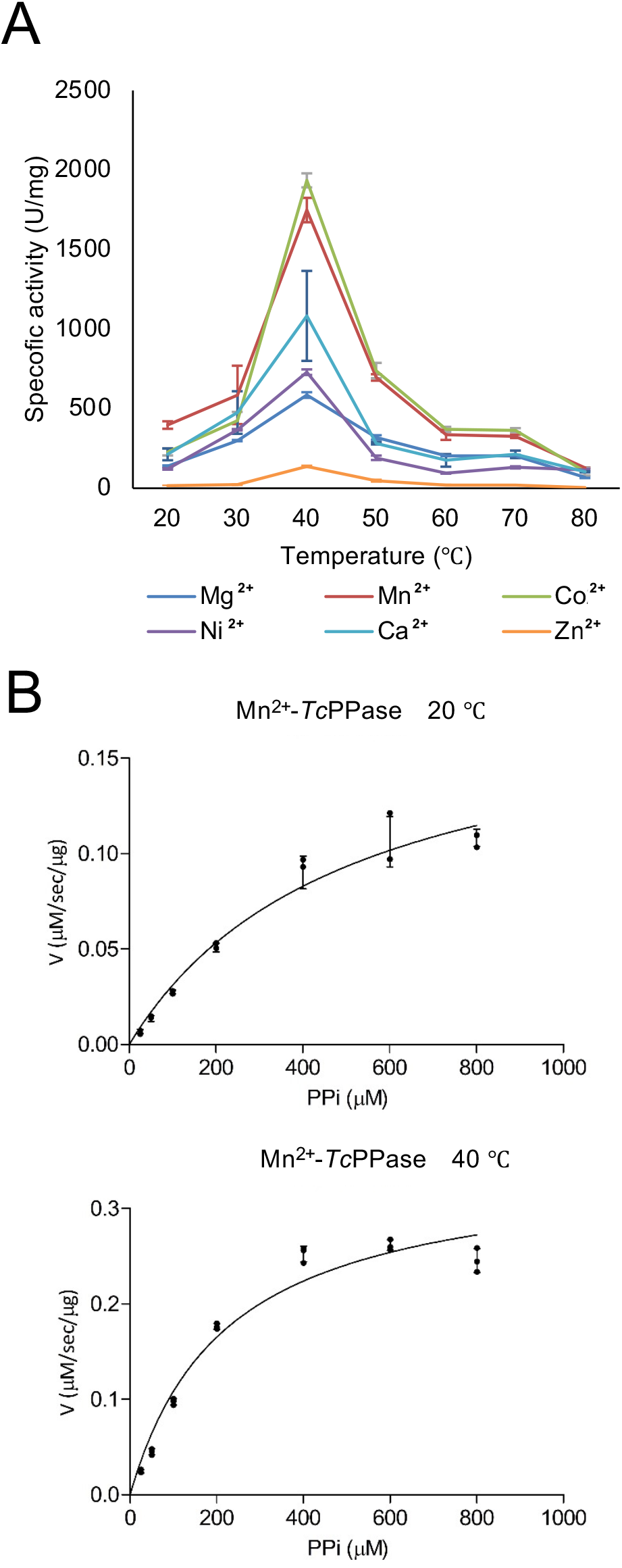
(A) Temperature dependence of *Thermodesulfobacterium commune* family II pyrophosphatase (*Tc*PPase) activity in the presence of different divalent metal ions under identical assay conditions. Data are represented as the mean ± standard deviation (SD) of three independent experiments. Different colors indicate the specific activities of Mg^2+^-(blue), Mn^2+^-(red), Co^2+^-(green), Ni^2+^-(purple), Ca^2+^-(cyan), and Zn^2+^-*Tc*PPase (orange). (B) Kinetic assays of Mn^2+^-*Tc*PPase at 20 °C (left) and 40 °C (right). Results are expressed as the mean ± SD of three independent experiments.

### Kinetic properties

Kinetic measurements of Mn^2+^-*Tc*PPase revealed distinct catalytic behavior at 40 °C compared to that at 20 °C. At 40 °C, *K*_*m*_ (221 ± 35 µM) was approximately half of that at 20 °C (500 ± 93 µM), and *k*_*cat*_ (122,000 ± 7,000 s^-1^) was about twice as high as that at 20 °C (65,500 ± 6,100 s^-1^; Fig. 1B; Table 1). These changes resulted in a higher *k*_*cat*_/*K*_*m*_ ratio at 40 °C, reflecting the enhanced substrate affinity and catalytic turnover relative to that at 20 °C. Compared to the reported values for mesophilic family II PPases at near-ambient temperatures [2,15,22,23,25,26], Mn^2+^-*Tc*PPase exhibited higher *k*_*cat*_ values, even at 20 °C.

**Table 1.** Kinetic parameters for pyrophosphate hydrolysis by Mn^2+^-*Thermodesulfobacterium commune* family II pyrophosphatase (*Tc*PPase). Data are represented as the mean ± standard deviation (SD) of three independent experiments.

| Temperature (°C) | 20 | 40 |
| --- | --- | --- |
| $k_{cat}$ (s<sup>-1</sup>) | 65,500 ± 6,100 | 122,000 ± 7,000 |
| $K_m$ (μM) | 500 ± 93 | 221 ± 35 |
| $k_{cat}/K_m$ (s<sup>-1</sup>μM<sup>-1</sup>) | 131 | 554 |

Considering the general temperature dependence of chemical reactions, where reaction rates typically increase 4–9-fold at every 20 °C rise in temperature [27], the observed 2-fold increase in *k*_*cat*_ for Mn^2+^-*Tc*PPase was modest. This suggests that *Tc*PPase evolved to optimize its catalytic efficiency within its functional temperature range by balancing its thermal stability and catalytic activity instead of significantly enhancing the turnover number. Interestingly, although binding to PPi in the active site involves hydrogen bonds and ionic interactions, which typically weaken as the temperature increases due to increased molecular motion [28], *Tc*PPase prioritized substrate affinity at elevated temperatures. This behavior suggests that its rigid structural features optimized for high-temperature stability possibly facilitate the structural changes necessary for substrate binding at elevated temperatures.

### Crystal structure and active site

Crystal structure of Mn^2+^-*Tc*PPase was determined at 2.06 Å resolution (Table S1). Despite attempts at crystallization in the presence of PNP, the obtained structure corresponded to the apo form. The enzyme consisted of an N-terminal DHH domain and a C-terminal DHHA2 domain and adopted an open conformation similar to *Bs*PPase (PDB: 1K23) [29] (Fig. 2A; Fig. S1). Root mean square deviation values of the aligned Cα atoms in the N-terminal DHH and C-terminal DHHA2 domains were 0.753 and 0.711 Å, respectively. Structural comparisons revealed that Mn^2+^-*Tc*PPase was closely aligned with other family II PPases with fully conserved catalytic residues (Fig. S2).

**Fig. 2.**
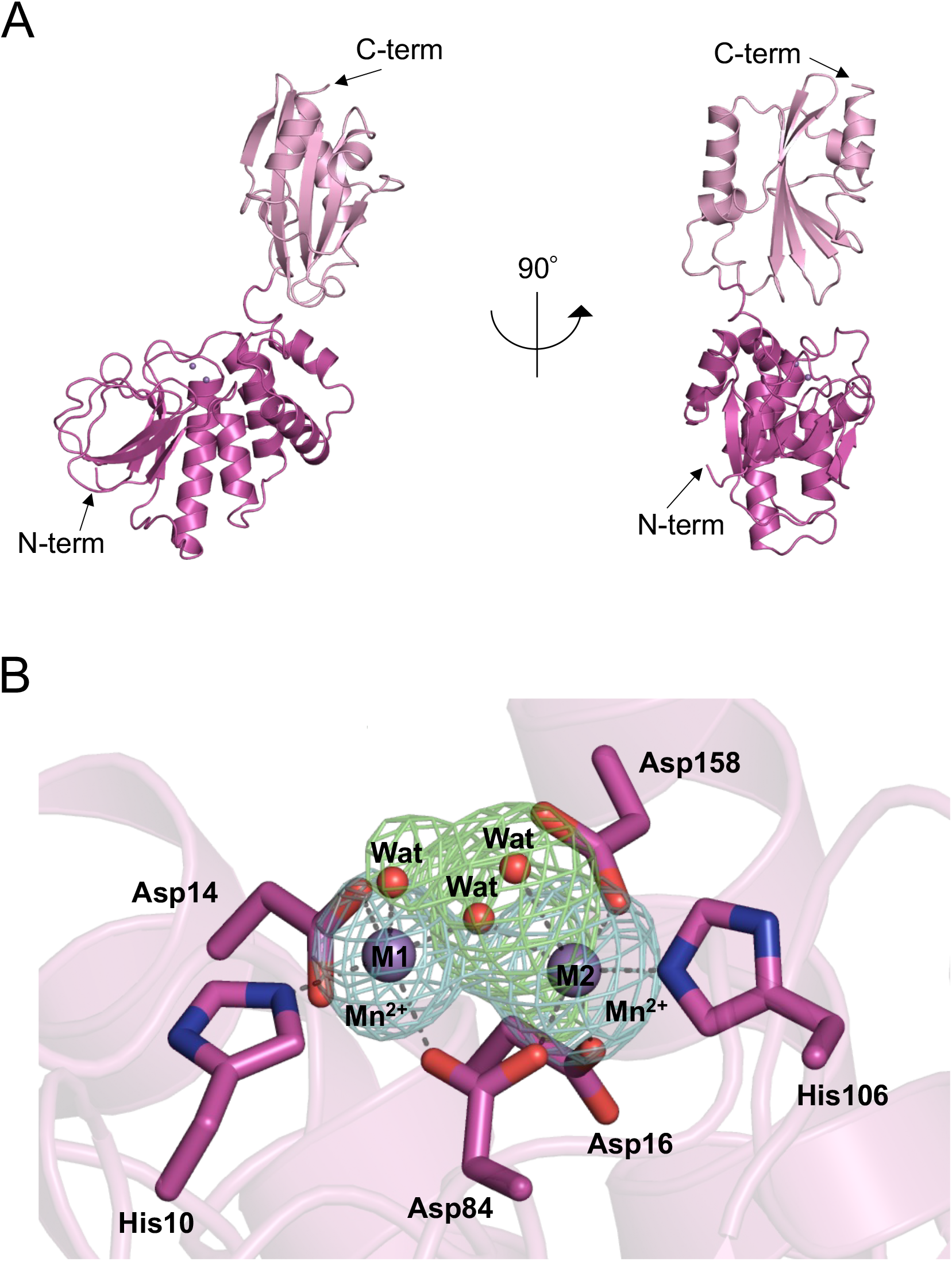
(A) Cartoon representation of the overall structure of Mn^2+^-*Tc*PPase. Right panel shows the molecule after a 90° rotation relative to the left panel. N- and C-termini (N-term and C-term, respectively) are indicated. N-terminal DHH and C-terminal DHHA2 domains are indicated in magenta and light pink, respectively. (B) Metal-binding site of Mn^2+^-*Tc*PPase with the corresponding polder maps. Key residues are shown as magenta sticks, with oxygen and nitrogen indicated in red and blue, respectively. Mn^2+^ ions are shown as deep purple spheres. Polder maps were contoured at 8s for Mn^2+^ ions (light blue) and 4s for water molecules (light green). Mn^2+^ ion coordination bonds are indicated by gray dashed lines.

M1 and M2 metal-binding sites in the active site showed coordination geometries typical of family II PPases [30,31], with M1 exhibiting a five-coordinate bipyramidal geometry, whereas M2 exhibiting a six-coordinate octahedral geometry (Fig. 2B). M1 site was coordinated to one nitrogen atom of histidine (His10), two oxygen atoms of aspartates (Asp14 and Asp84), and two oxygen atoms of water (metal-bridging and coordinated water). M2 site was coordinated to one nitrogen atom of histidine (His106), three oxygen atoms of aspartates (Asp16, Asp84, and Asp158), and two oxygen atoms of water (metal-bridging and coordinated water). Intermetal distance between the bridging water and M1 and M2 sites was 2.1 Å (Fig. 2B).

Structural superposition of the N-terminal DHH domain with the closed conformation of Mg^2+^-*Sh*PPase (PDB: 6LL8) [14] revealed a Cα interatomic distance of 49.2 Å between the C-terminal residue (Leu320) of Mn^2+^-*Tc*PPase and its counterpart in Mg^2+^-*Sh*PPase (Fig. S3). Dihedral angle formed by N-terminal His2 (N-terminal DHH domain), Asp200 (hinge), and C-terminal Leu320 (C-terminal DHHA2 domain) in Mn^2+^-*Tc*PPase and Mg^2+^-*Sh*PPase was 11.2°, indicating slight torsional differences between the two conformations (Fig. S3).

Closed-form Mn^2+^-*Tc*PPase model was derived by the independent superposition of the N-terminal DHH and C-terminal DHHA2 domains of Mn^2+^-*Tc*PPase and Mg^2+^-*Sh*PPase. This structural comparison indicated that substrate binding induced a conformational shift to a closed conformation, which repositioned Asp158 and Lys306 to stabilize the substrate and metal clusters for catalysis (Fig. 3). In the N-terminal DHH domain, residues involved in metal binding (Asp14, Asp16, Asp84, His106, and Asp158) aligned well with their counterparts in Mg^2+^-*Sh*PPase; however, Asp158 exhibited a conformational shift possibly induced by substrate binding to optimize the trimetal coordination essential for catalysis [14,30]. In the C-terminal DHHA2 domain, residues associated with PNP substrate binding (His107, Lys306, Lys215, and Arg305) were also aligned with those in Mg^2+^-*Sh*PPase. Notably, Lys306 underwent a conformational shift to accommodate PNP binding, thereby resulting in a closed conformation.

**Fig. 3.**
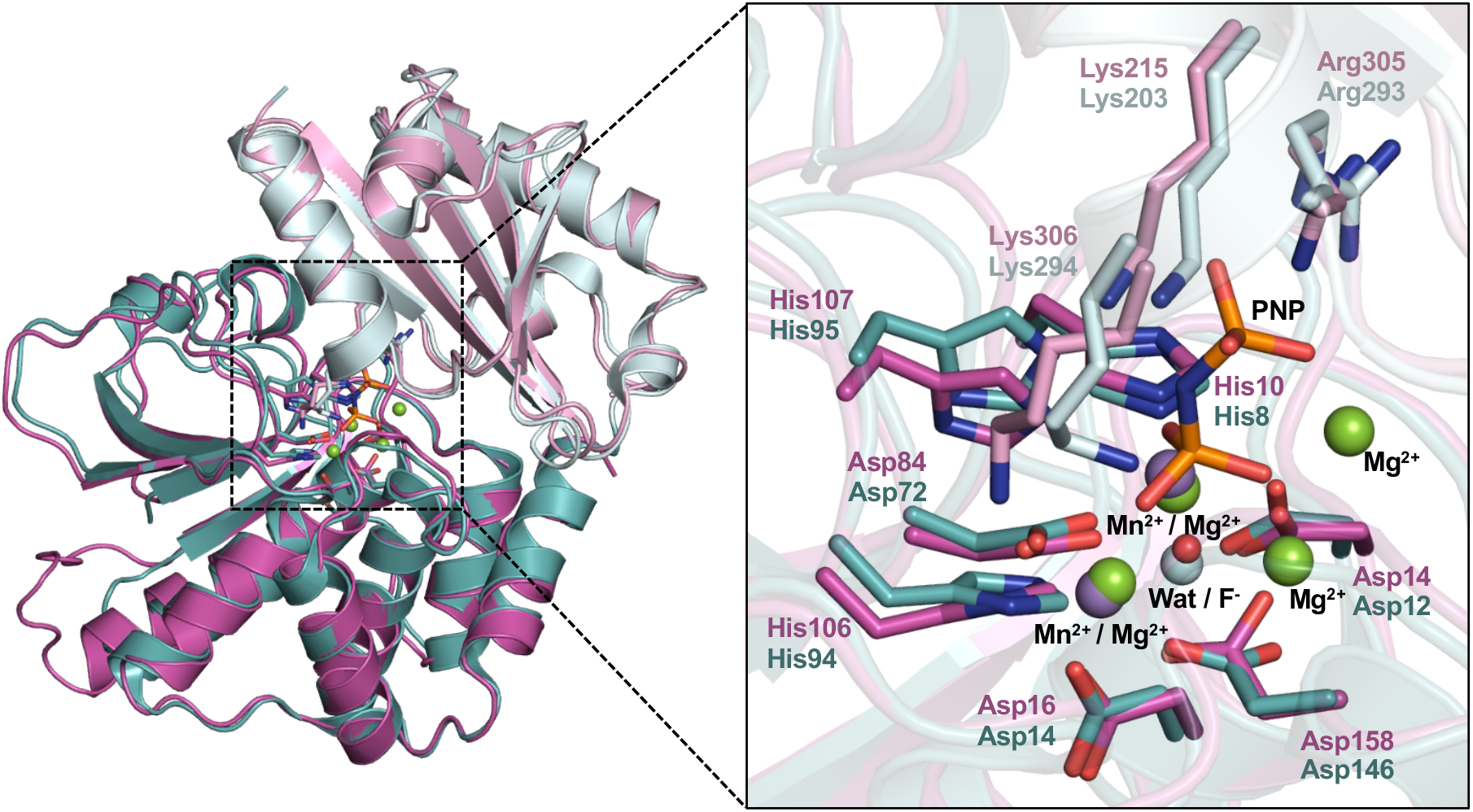
Closed-form Mn^2+^-*Tc*PPase model obtained via superposition with Mg^2+^-*Sh*PPase (PDB: 6LL8). N-terminal DHH and C-terminal DHHA2 domains were independently aligned based on the Cα atoms to achieve the closed conformation. N-terminal DHH domains of Mn^2+^-*Tc*PPase and Mg^2+^-*Sh*PPase are shown in magenta and cyan, respectively, and their C-terminal DHHA2 domains are indicated in light pink and light blue, respectively. Oxygen, nitrogen, and phosphorus atoms are indicated in red, blue, and orange, respectively. Mn^2+^ and Mg^2+^ ions are shown as deep purple and green spheres, respectively. Right panel shows a close-up view of the enzyme active sites, including the substrate analog (PNP) derived from the enzyme–substrate (ES) complex of Mg^2+^-*Sh*PPase.

### Comparative structural analyses for thermal adaptation

Comparative structural and surface analyses of the apo forms of *Tc*PPase, mesophilic *Bs*PPase (PDB: 1K23), and psychrophilic *Sh*PPase (PDB: 6LL7; Table 2) revealed that *Tc*PPase exhibited a larger hydrophobic surface area (8396 Å^2^) than *Bs*PPase (7213 Å^2^) and *Sh*PPase (7252 Å^2^) but a smaller hydrophilic surface area (4996 Å^2^) than *Bs*PPase (6107 Å^2^). Although a reduction in the hydrophobic surface area is often linked to increased protein stability [11,32], our results indicated that *Tc*PPase used an alternative stabilization strategy. The accessible surface areas of the enzymes were similar when accounting for their sizes, with *Tc*PPase at 14,105 Å^2^, *Bs*PPase at 13,944 Å^2^, and *Sh*PPase at 12,558 Å^2^. However, *Tc*PPase contained a higher proportion of buried hydrophobic residues (18%), corresponding to 57 buried hydrophobic residues out of the total 320 residues in the enzyme. Comparatively, *Bs*PPase and *Sh*PPase exhibited lower residue proportions of 16% (48 out of 309 residues) and 15% (47 out of 309 residues), respectively, indicating the enhanced hydrophobic interactions within the *Tc*PPase core, which possibly stabilize the core and strengthen the structural rigidity under high-temperature conditions [11,33,34].

**Table 2.** Structural parameters related to the protein stabilities of TcPPase, Bacillus subtilis family II PPase (BsPPase), and ShPPase. Apo-form structures were used for the comparison of TcPPase, BsPPase (Protein Data Bank [PDB]: 1K23), and ShPPase (PDB: 6ll7).

| Property | Thermophilic | Mesophilic | Psychrophilic |
| --- | --- | --- | --- |
|  | *TcPPase* | *BsPPase* | *ShPPase* |
| Number of residues | 320 | 309 | 309 |
| Molecular mass (kDa) | 35 | 34 | 34 |
| Hydrophobic surface area (Å<sup>2</sup>) | 8396 | 7213 | 7252 |
| Hydrophilic surface area (Å<sup>2</sup>) | 4996 | 6107 | 4721 |
| Accesible surface area (Å<sup>2</sup>) | 14105 | 13944 | 12558 |
| Buried hydrophobic residues content (%) | 18 | 16 | 15 |
| Proline residues content (%) | 4 | 2 | 4 |
| Helix content (%) | 39 | 40 | 39 |
| Strand content (%) | 21 | 20 | 20 |
| H-bonds per residue (cutoff $\leq 3.4\text{\AA}$) | 2.4 | 2.3 | 2.3 |
| Salt bridges per residue (cutoff $\leq 3.2\text{\AA}$) | 0.26 | 0.20 | 0.16 |

*Tc*PPase contains higher proline residue content (4%) than *Bs*PPase (2%), possibly contributing to its tighter protein conformation by reducing the backbone flexibility, thereby improving the thermal stability [34,35]. *Tc*PPase also exhibited a denser hydrogen bond network with 2.4 hydrogen bonds/residue compared to the 2.3 hydrogen bonds/residue in *Bs*PPase and 2.3 hydrogen bonds/residue in *Sh*PPase. Furthermore, *Tc*PPase had more salt bridges/residue (0.26) than *Bs*PPase (0.20) and *Sh*PPase (0.16), contributing to its enhanced structural stability via electrostatic interactions and tighter packing.

Notably, secondary structures, including helices (39% for *Tc*PPase, 40% for *Bs*PPase, and 39% for *Sh*PPase) and strand ratios (21% for *Tc*PPase, 21% for *Bs*PPase, and 20% for *Sh*PPase), were similar for these enzymes. This suggests that thermal adaptation in *Tc*PPase is driven primarily by intramolecular interactions, such as hydrophobic packing, hydrogen bonding, and salt bridge formation, rather than by large-scale secondary structure rearrangement.

## Discussion

This study is the first to report the successful expression, purification, and structural characterization of thermophilic family II PPases. *Tc*PPase required both the N-terminal DHH and C-terminal DHHA2 domains for its catalytic activity, similar to the domain interdependence observed in other family II PPases [14,36]. Crystal structure of Mn^2+^-*Tc*PPase, along with its kinetic and comparative analysis data, revealed that its thermal adaptation involved controlling the substrate affinity and turnover at elevated temperatures and reinforcing structural integrity via extensive hydrophobic interactions, increased proline content, dense hydrogen bonding, and additional salt bridges.

Comparative structural analysis of psychrophilic *Sh*PPase revealed its distinct environmental adaptation process. As shown in Table 2, *Sh*PPase possessed a lower proportion of buried hydrophobic residues (15%) compared to *Tc*PPase (18%) and *Bs*PPase (16%), along with the lowest hydrogen bond density (2.3/residue) and salt bridge density (0.16/residue) among the analyzed enzymes. These features contributed to its less rigid but more dynamic structure, enabling *Sh*PPase to maintain its structural flexibility and catalytic efficiency in cold environments [23,37]. The different adaptation strategies of *Tc*PPase and *Sh*PPase highlight the structural versatility of family II PPases under diverse thermal conditions.

Thermophilic *Tc*PPase uses its structural features to stabilize and tighten its structure at elevated temperatures, whereas psychrophilic *Sh*PPase relies on its structural flexibility to maintain activity at low temperatures. These strategies optimize the enzyme binding efficiency and catalytic performance at optimal temperatures. Interestingly, despite these structural differences, secondary structure compositions of *Tc*PPase, *Bs*PPase, and *Sh*PPase were remarkably similar. This suggests that thermal adaptation in family II PPases is primarily driven by the fine-tuning of intramolecular interactions within a conserved structural framework instead of the significant modification of the overall secondary structure.

In conclusion, our findings enhance the understanding of the diverse strategies used by family II PPases to adapt to extreme environmental conditions. Additionally, this study provides a foundation for thermostable enzyme engineering for various industrial and biotechnological applications requiring robust enzyme performance under harsh conditions.

## Acknowledgements

The synchrotron radiation experiments were performed at SPring-8 (proposal nos. 2021B2553 and 2022A2553). We thank the Center for Advanced Instrumental and Educational Support of the Faculty of Agriculture (Kyushu University) for technical support.

## Funding

This work was supported by a Research Fellowship for Young Scientists from the Japan Society for the Promotion of Science (JSPS) (JP23KJ1741 to S. M.).

## Abbreviations

PPases: inorganic pyrophosphatases
*Tc*PPase: *Thermodesulfobacterium commune* family II PPase
PPi: inorganic pyrophosphate
Pi: phosphate
PNP: imidodiphosphate sodium salt
*Bs*PPase: *Bacillus subtilis* family II PPase
*Sh*PPase: *Shewanella* sp. AS-11 family II PPase

---

**Fig. S1.**
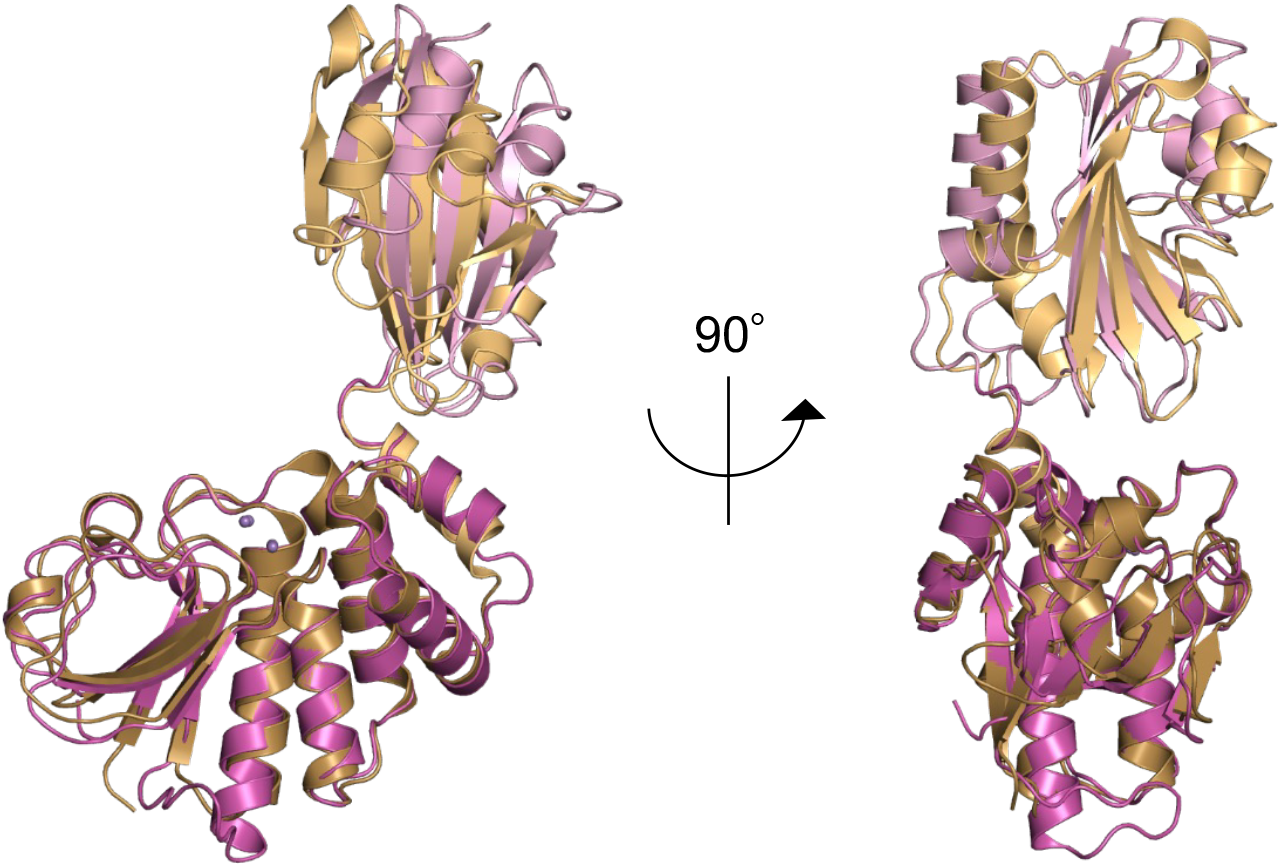
Superposition of Mn^2+^-*Tc*PPase and Mn^2+^-*Bs*PPase (PDB ID: 1K23) structures. Both structures are shown in cartoon representation. The N-terminal DHH domains of Mn^2+^-*Tc*PPase and Mn^2+^-*Bs*PPase are colored magenta and tan, respectively, and their C-terminal DHHA2 domains are light pink and beige. Mn^2^+ ions are depicted as deep purple spheres. The right panel shows the structure rotated by 90° relative to the left panel.

**Fig. S2.**
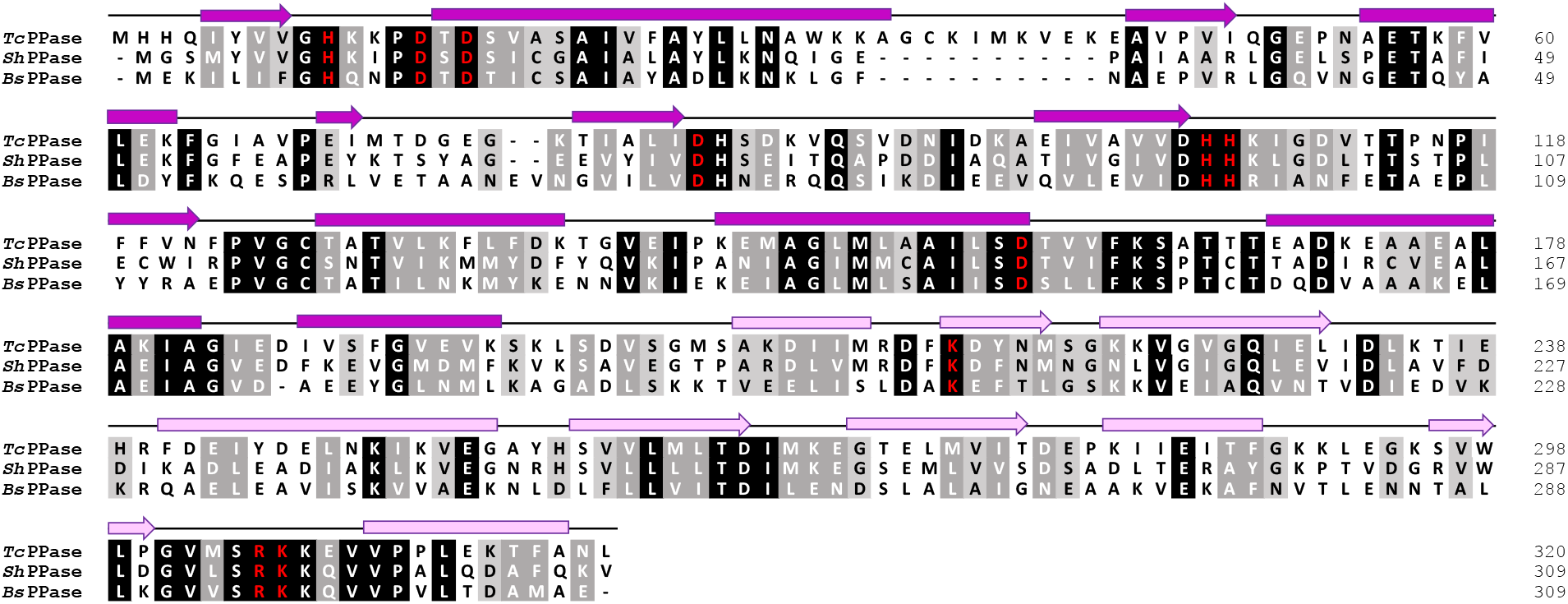
Multiple sequence alignment of *Tc*PPase, *Sh*PPase, and *Bs*PPase, shown together with the secondary structure elements of *Tc*PPase. Alpha-helices and β-strands are indicated by boxes and arrows, respectively. Conserved catalytic residues are highlighted in red. Background shading intensity reflects the degree of sequence conservation, with darker shading indicating higher conservation.

**Fig. S3.**
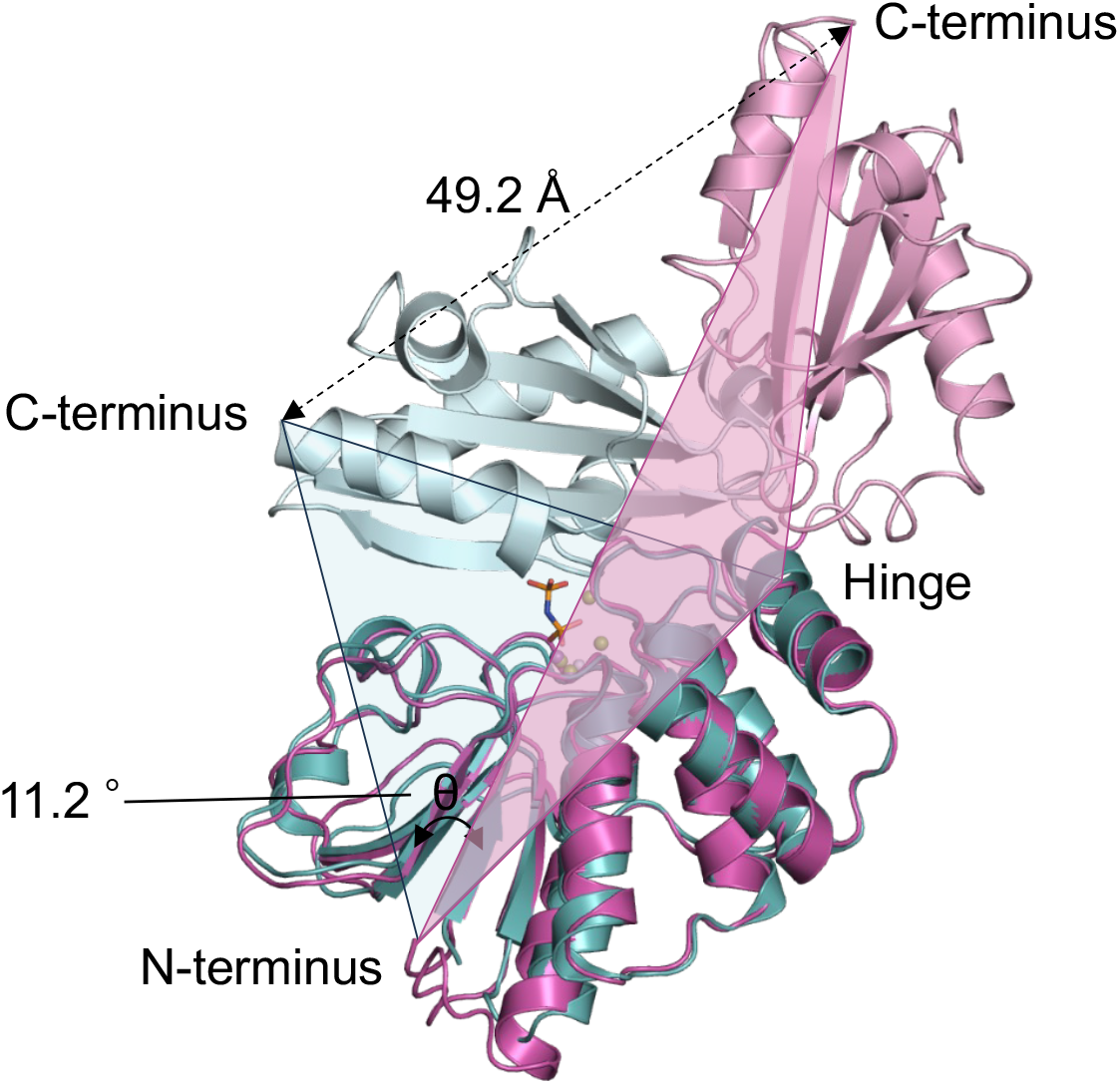
Superposition of the N-terminal DHH domains in Mn^2+^-*Tc*PPase and Mg^2+^-*Sh*PPase (PDB ID: 6ll8) structures. Both structures are displayed in cartoon representation. The N-terminal DHH domains of Mn^2^+ -*Tc*PPase and Mg^2^+-*Sh*PPase are colored magenta and cyan, respectively, while their C-terminal DHHA2 domains are shown in light pink and light blue. Mn^2^+ ions and Mg^2^+ions are depicted as deep purple and green spheres, respectively. The substrate analog PNP in Mg^2^+-*Sh*PPase is represented as sticks. The N-terminus-Hinge-C-terminus planes in Mn^2^+ -*Tc*PPase and Mg^2^+-*Sh*PPase are highlighted in magenta and cyan, respectively.

**Table S1.** Crystallographic data collection and refinement statistics.

|  |  |
| --- | --- |
| **Data collection** |  |
| Space group | P 6<sub>1</sub> 2 2 |
| Cell dimensions |  |
| *a*, *b*, *c* (Å) | 66.07 66.07 451.2 |
| α, β, γ (°) | 90 90 120 |
| Resolution (Å) | 45.54 – 2.08<br>(2.21 – 2.08)* |
| *R*<sub>means</sub> | 6.5 (191.3) |
| CC1/2 | 99.8 (61.6) |
| *I* / σ*I* | 37.72 (1.03) |
| Completeness | 99.1 (100.0) |
| (%) |  |
| Redundancy | 17.01 (21.62) |
| **Refinement** |  |
| Resolution (Å) | 45.54 – 2.08 |
| No. reflections | 36785 |
| *R*<sub>work</sub> / *R*<sub>free</sub> | 18.8 / 22.0 |
| No. atoms |  |
| Protein | 2613 |
| Ligand | 33 |
| *B*-factors |  |
| Protein | 71.5 |
| Ligand | 108.1 |
| R.m.s. deviations |  |
| Bond lengths | 0.003 |
| (Å) | 0.53 |
| Bond angles |  |
| (°) |  |
| Ramachandran |  |
| plot | 95.9 |
| Favored (%) | 4.1 |
| Allowed (%) | 0 |
| Disallowed |  |
| (%) |  |

\*Values in parentheses are for highest-resolution shell.

## Notes

### Competing Interest Statement

The authors have declared no competing interest.

